# Reconfiguration of directed functional connectivity among triple networks with aging: Considering the role of thalamo-cortical interactions

**DOI:** 10.1101/827451

**Authors:** Moumita Das, Vanshika Singh, Lucina Uddin, Arpan Banerjee, Dipanjan Roy

**Affiliations:** Cognitive Brain Dynamics Lab, National Brain Research Centre, NH 8, Manesar, Gurgaon 122052, India; Department of Psychology, University of Miami, Coral Gables, FL USA

**Keywords:** **Healthy** Aging, Directed Functional Connectivity, Multivariate Granger Causality, Salience Network, Thalamus, Weighted Net Causal Outflow

## Abstract

The human brain undergoes significant structural and functional changes across the lifespan. Our current understanding of the underlying causal relationships of dynamical changes in functional connectivity with age is limited. On average, functional connectivity *within* resting-state networks (RSNs) weakens in magnitude, while connections *between* RSNs tend to increase with age. Recent studies show that effective connectivity within and between large scale resting-state functional networks changes over the healthy lifespan. The vast majority of previous studies have focused primarily on characterizing cortical networks, with little work exploring the influence of subcortical nodes such as the thalamus on large-scale network interactions across the lifespan. Using directed connectivity and weighted net causal outflow measures applied to resting-state fMRI data, we examine the age-related changes in both cortical and thalamocortical causal interactions within and between RSNs. The three core neurocognitive networks from the triple network theory (default mode: DMN, salience: SN, and central executive: CEN) were identified independently using ICA and spatial matching of hub regions with these important RSNs previously reported in the literature. Multivariate granger causal analysis (GCA) was performed to test for directional connectivity and weighted causal outflow between selected nodes of RSNs accounting for thalamo-cortical interactions. Firstly, we observe that within-network causal connections become progressively weaker with age, and network dynamics are substantially reconfigured via strong thalamic drive particularly in the young group. Our findings manifest stronger between-network directional connectivity, which is further strongly mediated by the SN in flexible co-ordination with the CEN, and DMN in the old group compared with the young group. Hence, causal within- and between- triple network connectivity largely reflects age-associated effects of resting-state functional connectivity. Thalamo-cortical causality effects on the triple networks with age were next explored. We discovered that left and right thalamus exhibit substantial interactions with the triple networks and play a crucial role in the reconfiguration of directed connections and within network causal outflow. The SN displayed directed functional connectivity in strongly driving both the CEN and DMN to a greater extent in the older group. Notably, these results were largely replicated on an independent dataset of matched young and old individuals. Our findings based on directed functional connectivity and weighted causal outflow measures strengthen the hypothesis that balancing within and between network connectivity is perhaps critical for the preservation and flexibility of cognitive functioning with aging.

## 1. Introduction

In the last decade, there has been an enormous interest in studying the coordinated activity in distributed brain areas when the human being is engaged in internally driven tasks, such as meandering through self-referential thoughts while seemingly at rest, or more specifically not engaged in a state of goal directed action and perception (Raichle, 2009; (Bressler & Menon, 2010); (Deco, Jirsa, & McIntosh, 2011)). The functional organization (Bressler Menon, 2010) that dominates the landscape of both resting state and task activity has been broadly classified into three networks based on the correlation patterns estimated from BOLD time series signals. These networks are known as the default mode network (DMN)/medial frontoparietal network, salience network (SN)/midcingulo-insular network, and central executive network (CEN)/lateral frontoparietal network ((Menon 2011) (Uddin, Yeo, & Spreng, 2019)). The DMN comprises posterior cingulate cortex (PCC) and medial prefrontal cortex (MPFC) and is implicated in self-referential mental activities (Buckner, Andrews-Hanna, & Schacter, 2008), (Andrews-Hanna, Reidler, Sepulcre, Poulin, & Buckner, 2010), (Raichle, 2015)(Uddin, Iacoboni, Lange, Keenan, 2007, TICS)). The CEN comprises rostral and caudal bilateral middle frontal cortex (MFC), bilateral superior Parietal Cortex (SPC) and is implicated in decision making and executive functions (Corbetta & Shulman, 2002); (Fox, Corbetta, Snyder, Vincent, & Raichle, 2006)). The SN, which comprises bilateral anterior insula (AI) and caudal, rostral bilateral anterior cingulate cortices (ACC) is important for detection and mapping of salient inputs and routing these inputs to control areas for mediating cognitive control ((Menon & Uddin, 2010); (Uddin, 2015)).

Interestingly, modification of interconnections within and between the DMN, CEN, SN is altered in a large number of psychiatric and neurological disorders, for instance, Alzheimer’s disease, autism spectrum disorder (ASD), attention deficit/hyperactivity disorder (ADHD), psychosis and depression (Woodward & Cascio, 2015);(Abi-Dargham & Horga, 2016)). The SN, in addition to detecting salient stimuli, plays important role in switching between the DMN and CEN in task conditions as well as in the resting state ((Devarajan, Levitin, & Menon, 2008) (Uddin et al., 2011)). Thus, it mediates controlled switching between internal mental processes and external stimulus-driven cognitive and affective processes. During the performance of many tasks, correlation among the nodes of SN in tandem with CEN increases, while the corresponding correlation among the DMN nodes decreases.

An important caveat to all the resting brain network studies are that they mostly concentrate on cortical nodes and ignore the crucial influence of thalamocortical interactions on whole-brain network dynamics. The thalamus, a centrally located relay station for transmitting information throughout the brain, participates in communication with many associative brain regions and involves global multifunctional pathways. Incorporating 10,449 meta-studies, Hwang, Bertolero, Liu, & D’Esposito, 2017 recently showed that the thalamus is engaged in multiple cognitive functions and is a critical integrative hub for functional brain networks. Cross-sectional studies of normal aging have also reported smaller thalamic volumes in older than younger adults (Cherubini, Péran, Caltagirone, Sabatini, & Spalletta, 2009, Hughes et al., 2012;). However, exactly how normal aging affects thalamic interconnections with other brain networks and its implication in cognitive changes are not completely understood, and thus warrant further investigation. From a methodological standpoint, if the thalamus acts as a common source to cortical inputs and also receives feedback from the cortex as proposed by theories such as thalamocortical dysryhtmia ((Llinás, Ribary, Jeanmonod, Kronberg, & Mitra, 1999),(Vanneste, Song, & De Ridder, 2018)), a causality analysis that ignores thalamocortical contribution to brain dynamics is of very limited scope, and possibly paints an inaccurate account of the underlying complexity. Thus, the primary goal of this study is to re-evaluate dynamic reconfiguration patterns in the triple networks by accounting for thalamo-cortical causal interactions and directed functional connectivity associated with aging. Our findings may be a first step for an understanding more general interactions between triple networks and subcortical structures to facilitate bidirectional causal information processing at the functional level.

## 2. Material and Methods

### 2.1 Participants

25 young and 24 elderly individuals participated in this study after providing written informed consent. The young group ranged in age from 18-33 years (mean age=25.7±4 yrs, 13 female) and the elderly group ranged in age from 55-80 years (mean age=67.99 ±9years, 18 female). The study was performed under the compliance of laws and guidelines approved by the ethics committee of Charité University, Berlin. In the replication cohort, we identified a group of 24 young (Age: 24.58 years; Range :18-31years; 11 female) and 25 elderly participants (Age: 64.8 years; Range : 50-81 years; 18 female) from the publicly available Cambridge Aging Neuro-science dataset (https://camcan-archive.mrc-cbu.cam.ac.uk//dataaccess/) who were similar in mean age and gender distribution to the Berlin dataset (see Table 2 for demographic details)

### 2.2 Data Acquisition

Resting state fMRI as well as diffusion weighted (dw) MRI data were collected from 49 healthy participants at the Charité University Berlin, Germany (Schirner, Rothmeier, Jirsa, McIntosh, & Ritter, 2015). Each fMRI dataset amounts to 661 time points recorded at TR=2s, i.e. about 22 minutes. In the same session, EEG was also recorded, but these data are not used for our current analysis. No other controlled task was performed. Resting state BOLD activity was recorded while subjects were asked to stay awake with their eyes closed, using a 3T Siemens Trim Trio scanner and a 12 channel Siemens head coil (voxel size).

### 2.3 Data Analysis

#### 2.3.1 rs-fMRI preprocessing

The major pre-processing steps applied to T1 anatomical images were skull stripping, removal of non-brain tissue, brain mask generation, cortical reconstruction, motion correction, intensity normalization, white matter (WM) and subcortical segmentation, cortical tessellation generating GM–WM and GM–pia interface surface-triangulations and probabilistic atlas-based cortical and subcortical parcellation. The parcellation used in this study is Desikan–Killiany parcellation (Desikan et al., 2006) which consists of 68 ROIs with 34 ROIs in each hemisphere and 32 subcortical 32 subcortical regions. For our present analysis along with 68 cortical regions, two thalamic regions, left and right thalamus were selected based on this parcellation. FC matrices generated from each subject’s MRI data are averaged element-wise to obtain an averaged FC matrices for the younger and elderly cohorts. The regional resting-state fMRI time series was computed for each of the regions-of-interest (ROIs, defined by the Desikan-Killiany atlas (Desikan et al., 2006) as implemented in FreeSurfer) by averaging all the voxels within each region at each time point in the time series.

These regional time series were temporally filtered using a bandpass filter (0.01 Hz < f < 0.08 Hz). The empirical BOLD time series signals from ROIs used in this paper for the estimation of directed connectivity and weighted net causal flow were generated by using an automated pipeline as described in detail elsewhere ((Schirner, Rothmeier, Jirsa, McIntosh, & Ritter, 2015)).

#### 2.3.2 Selection and extraction of triple networks

To identify RSN activity, a spatial Group ICA decomposition was performed for the fMRI data of all subjects using FSL MELODIC ((Beckmann & Smith, 2004)) (MELODIC v4.0; FMRIB Oxford University, UK) with the following parameters: high pass filter cut off: 100 s, MCFLIRT motion correction, BET brain extraction, spatial smoothing, normalization to MNI152, temporal concatenation, dimensionality restriction to 30 output components. ICs that correspond to RSNs were automatically identified by spatial correlation with the 9 out of the 10 well-matched pairs of networks of the 29,671-subject Brain Map activation database as described in (Smith et al., 2009) (excluding the cerebellum network). Subsequently, the three key intrinsic networks were identified by spatially matching with pre-existing templates following widely accepted seven networks resting state parcellation proposed by Buckner and colleagues (Buckner, Krienen, Castellanos, Diaz, & Yeo, 2011). Each of the cortical regions were classified further down to seven parcellated resting networks. In our subsequent analysis, we considered three resting state networks, namely the DMN, SN, CEN (Node details are provided in **Table: 1**). We chose for our study Inferior parietal lobule (IPL), PCC and MOF as they constitute core part of DMN ((Andrews-Hanna et al., 2010); (Dixon et al., 2017)) consistently showed activation during mental rumination and self-related processing, mind wandering. The SN, which comprises the anterior insula and caudal, rostral anterior cingulate cortex, is important for detection of salience events and switching between other large-scale networks ((Menon & Uddin, 2010); (Uddin, 2015)). Bilateral rostral and caudal middle frontal gyrus (MFG), superior parietal lobule (SPL) were selected as nodes of CEN. The core DMN regions selected in this work consistently showed anticorrelation with SNs ((Fransson, 2005); (Uddin, Kelly, Biswal, Castellanos, & Milham, 2009); (Dixon et al., 2017)).

**Table 1:** Coordinates of selected nodes of three resting state networks according to Desikan-Killiany (DK) parcellation atlas

| Networks | Brain Regions | MNI coordinates<br>(x,y,z) |  |
| --- | --- | --- | --- |
|  |  | Left (l) | Right(r) |
| Salience Network | Insula (Insula) | (-41, 13, -6) | (43, 12, -6) |
|  | Caudal anterior cingulate cortex (CACC) | (-2,21,27) | (3, 21, 27) |
|  | Rostral anterior cingulate cortex (RACC) | (-2,39,6) | (4, 38,4) |
| Central Executive Network | Rostral middle frontal gyrus (RMFG) | (-34, 53, 17) | (43, 45, 21) |
|  | Caudal middle frontal gyrus (CMFG) | (-45, 18, 46) | (43, 14, 43) |
|  | Superior parietal lobule (SPL) | (-25, -62, 63) | (17, -65, 59) |
| Default Mode Network | Medial orbitofrontal (MOF) | (-4, 44, -14) | (7, 45, -13) |
|  | Inferior parietal lobule (IPL) | (-47, -70, 31) | (48, -67, 29) |
|  | Posterior cingulate cortex (PCC) | (-1, -18, 38) | (1, -16, 37) |

For within network analysis, we used the extracted time series for each of the six ROIs within each respective network. For between network analysis, we combined time series of the six ROIs using principal component analysis to create a single representative time series of each network.

#### 2.3.3 Directed Functional Connectivity

The main objective of the present study is to explore the directional changes and weighted causal outflow in the well-known triple resting state networks across the lifespan and in the presence of exogenous drive from bilateral thalamic. In order to estimate directional connectivity in the time domain and also in the frequency domain, we employed Multivariate Granger causality analysis that is particularly suitable for the exploratory nature of our study (Reid et al., 2019).

### 2.4 Multivariate Granger Causality Analysis

If we consider two variables X and Y that evolve over time, Y is said to G-cause X if the past values of Y contain information that helps predict the future of X over and above information in the past values of X alone (Granger, 1969; Geweke, 1982, 1984). This can be tested by constructing a univariate autoregression model of X and checking whether or not the lagged values of Y add any explanatory power to the model. However, neural signals from multiple brain regions are expected to show joint dependencies, thus necessitating the multi-variate analysis. For instance, let ***X_t_***, ***Y_t_***, ***Z_t_*** be the time series of 3 nodes in a network. If there are joint dependencies between ***X_t_***, ***Y_t_***, ***Z_t_*** and if we calculate unconditional granger causality between ***X*** and ***Y***, spurious causalities may occur due to the common dependency on ***Z***. Thus, to eliminate the possibility of spurious causalities between two time series, Multivariate Granger Causality analysis (GCA) was performed to assess the causal influence between nodes of the SN, CEN and DMN based on the methods described in ((Barrett, Barnett, & Seth, 2010)). In MVGC, spurious causalities are eliminated by conditioning out the common dependencies. Thus, to test the G-causality from ***Y*** to ***X***, one needs to consider the full and reduced regressions of the following form:

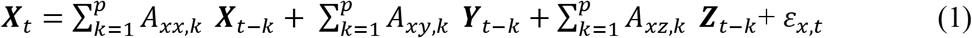

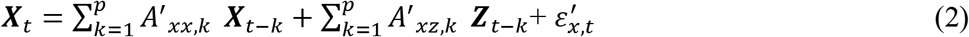

Here *p* is the model order, *A* is the regression coefficient, and *ε* the residuals.. In full regression (1), the dependence of X on the past values of Y, given its own past values and the past values of Z is incorporated in the coefficients *A_xy,k_*.. If there is no conditional dependence of X on the past values of Y, this coefficient becomes zero. This leads to the reduced regression equation (2). Therefore, the null hypothesis of zero causality is:

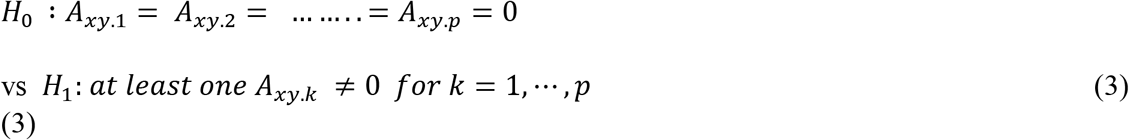

The G-causality value quantifies the degree to which the full regression model is a better candidate compared to the reduced regression to model ***X_t_***. An appropriate measure for the model comparison or prediction error in directional connectivity between selected pairs of nodes is the logarithm of the ratio of their likelihood values. The joint likelihood of the var model is 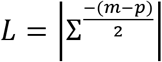 ((where |Σ| =the *generalised variance* of the model, *m* = total number of nodes, *p* = estimated model order). The conditional G -causality from Y to X in equation (1) and (2) is defined as:

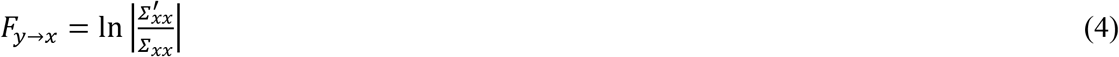

Where 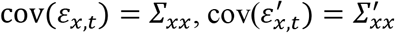 are the error variances of the full model and reduced model, respectively. Although three time series were considered in the above example, in general, vector autoregressive (VAR) modelling is employed for each time *t* deals with a *n*-dimensional vector space with column vector *U* represents a multivariate time series signal. Therefore, a *p*-th order VAR model can be represented using the similar notations as above in equations (1) and (2) as:

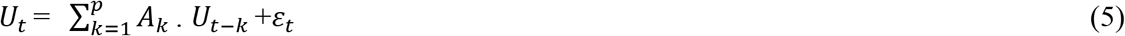

We employed the Multivariate Granger Causality toolbox (Barnett et al.(2014)) to perform the connectivity analysis. Joint likelihood *L* was calculated for each model order *p* up to the maximum model order. In practice, given empirical time series data, the model order *p* should be selected based on some theoretical criterion. The value of *p* should be sufficiently large to capture the predictive variation, but very large values of *p* are also not desirable, because they will induce over-fitting in the model. The order of the VAR model used for computation of the influence measure was selected using the Bayesian information criterion (BIC) (Barnett et al.(2014)). The model with lowest BIC was chosen. Corresponding VAR model parameters, VAR model coefficients, and covariance matrices were estimated for the estimated model order. Using the reverse solution of the Yule Walker equations (Barnett et al.(2014)), the autocovariance sequences were calculated. For Granger causal estimation, VAR parameters were calculated for both the full and reduced regressions. Granger causality value was calculated using (4). Significance was tested using nonparametric analysis and *F*-tests as employed in (Uddin et al. 2011, Barnett et al.(2014)). Empirical null distributions of the influence of one node on another based on *F*-values and their differences were estimated nonparametrically using bootstrapping and surrogate analysis with null hypothesis of no causal interactions between the brain regions. Since multiple causalities were tested simultaneously, false discovery rate (FDR) correction was used to adjust for multiple hypotheses. The main strength of MVGC is that using the multiple equivalent representation of a VAR model by regression parameters, the autocovariance sequence, one is able to compute G-causality with the simultaneous estimation of full and reduced regression. This increases the power of statistical tests as well as model estimation accuracy.

#### 2.4.1 Network Analysis

We estimated and quantified the following metrics to further characterize the networks in the young and elderly cohorts: (1) out-degree, number of causal outflow connections from a node in the network to any other node; (2) in-degree, number of causal in-flow connections to a node in the network from any other node; and (3) (out–in) degree, difference between out degree and in degree, a measure of the net causal outflow from a node. This permitted comparison with previously reported causal outflow measures in large-scale brain networks (Devarajan, Levitin, & Menon, 2008; Uddin et al., 2011).

We calculated the weighted net (Granger) causal flow. The main differences from previous work were that we employed here weighted net causal flow of a node, that was defined as the weighted (Out-In) degree instead of what was previously employed and described above. This decision was made to account for the fact that unweighted estimates actually may lead to incorrect inferences about causal structure. Weighted out-degree of a node is the sum of the strength (i.e., Granger causal indices) of significant causal connections from the node to all the other nodes in the network. Likewise, weighted in-degree of a node is the sum of the strength of significant causal inflow connections to a that node from remaining nodes in the network. For example, weighted net causal outflow of node X, say Δ*_x_* can be expressed using the following formula:

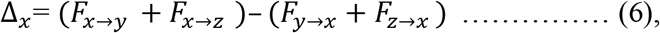

provide all *F* values are significant. Non-significant *F*-values should be instead replaced by zero. Similarly, one can calculate Δ*_y_*, Δ*_z_*. Though *F* values are positive, weighted net causal values can be positive as well as negative. Positive Δ for a particular node implies higher causal influence of that node on the other nodes of the network, whereas negative Δ signifies-that the particular node is more causally driven by other nodes of the network (in other words more inflow). These net causal outflow measures are used to characterise the hub of any causal flow network and redefine the role of hubs with age-associated alterations. In this definition, the node with the highest net causal outflow in a network is considered to be the hub of that particular network.

To compare the net flow of different nodes between young and old age groups we generated 100 samples of Granger causality index matrices via nonparametric bootstrap sampling. For the purpose of comparison, we constructed the distribution of weighted net causal outflow based on the significant causal connections (p<0.05, FDR corrected for multiple comparisons). The distribution of weighted net causal outflow was calculated for different nodes within and between networks, and Wilcoxon signed rank tests were performed to test the significant differences in the net causal outflows for the different age groups.

### 2.5 Patterns of age associated decrease within triple network causality influenced by thalamo-cortical causal interactions

Next, we ask how much of the within network reduction in directed connectivity and weighted causal flow (In-out degree) as observed is actually shaped up by declining cortical-thalamic causal connectivity with age. To this end, we included the thalamus as an additional node in our network analysis to track thalamo-cortical directed connectivity. Hence, assuming we have three variables: *X (cortical node 1), Y (cortical node 2)*, and *Z (subcortical node)*, and are interested in measuring information flow from *X* to *Y*. First, a “full” VAR model (Eq. 1) is estimated jointly for all three variables. This leads to a particular prediction/estimation error for each variable within the set. A second “reduced” VAR model is then estimated, which omits the potential cause (*X*, in the example above). This leads to a second set of prediction errors for each remaining variable. If the prediction error for *Y* is significantly smaller for the full regression (including *X*), as compared with the reduced regression (Eq. 2) (excluding *X*), then we say that *X* G-causes *Y*, conditioned on *Z*. (Note that we also have, from the same models, the G-causality from *X* to *Z* conditioned on *Y*). Technically, the magnitude of the G-causality is given by the ratio of the variance of the prediction-error terms for the reduced and full regressions (F values estimating prediction error).

For each of the three resting state networks, for within network analysis, we included the right and the left thalami as the seventh and the eighth nodes. Since the mechanism of the MVGC changes with the number of nodes, the causal strengths were not directly comparable between two networks having a different number of nodes. In other words, it is not meaningful to quantify the difference between causal strengths of the same RSN networks before and after the inclusion of the thalami. Instead of that, we focused our analysis on discovering changes in the pattern of causal connections and net causal outflows.

### 2.6 Code Accessibility

All code used for fMRI data analysis, causality estimation and generation of manuscript figures are freely available from https://github.com/dynamicdip/CBDL_Granger_causal_AGING_fMRI and also upon reasonable request from the corresponding author. The analysis code has been tested on both Linux and Mac OSX platforms.

## 3.0 Results

### 3.1 Age-associated within and between network reconfiguration in causal connectivity structure in triple networks

#### 3.1.1 Comparison of within network directed causal connectivity between network nodes in young versus elderly using multivariate GCA in CEN, SN and DMN

We used multivariate GCA (Barnett et al. (2014)) to investigate causal interactions between the six network nodes in the three well studied neurocognitive networks. In the CEN, we extracted time series from six nodes (see **figure 2**), namely lRMFG, rRMFG, lCMFG, rCMFG, lSPL, rSPL. The strength of the directed interaction from the rRMFG (driver node in a network) to the rCMFG (follower node in a network) was highest among all causal interactions for both the groups (**figure 3A, 3D**). There were no bidirectional interactions in the CEN. Interestingly, we did find rostro-caudal directed interactions in the CEN. Furthermore, there were substantial unidirectional drive from anterior regions of the CEN to the posterior regions along the anterior-posterior axis. We found significant directed connectivity (*p* < 0.01, FDR corrected) from the lRMFG to the lCMFG, and the lSPL for young individuals (restricted to a single hemisphere) (**figure 3D**). On the other hand, for the elderly group, significant directed connectivity were from the rRMFG to the lSPL (between two hemispheres) and rSPL (in addition to the rCMFG) (**figure 3A**). In contrast to DMN, a greater number of inter hemispheric connections were found in elderly individuals in the CEN (**figure 3A**).

**Figure 1.**
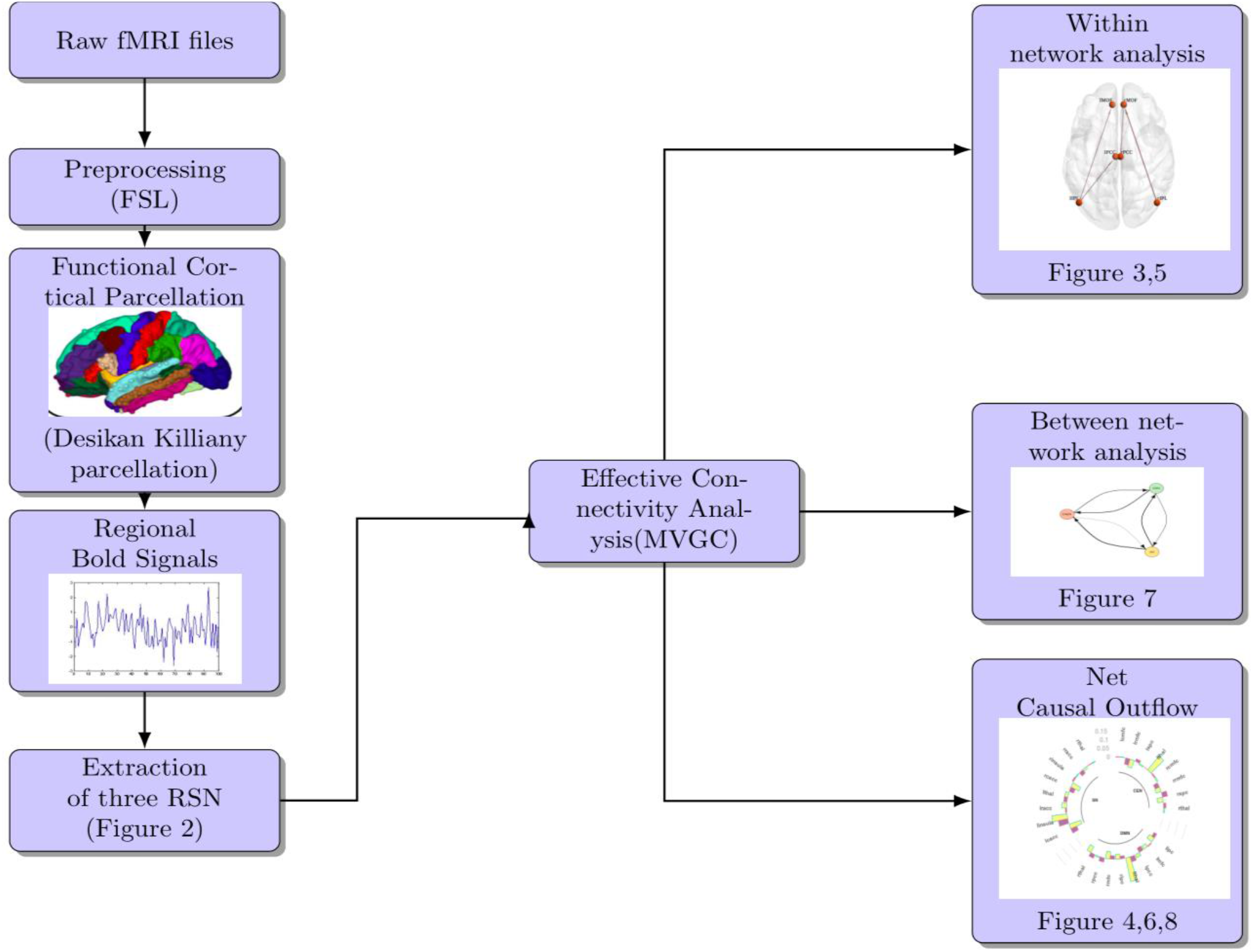
Flow chart describing major steps employed in our pipeline for estimation of causal connectivity and weighted outflow analysis using MVGC

**Figure 2.**
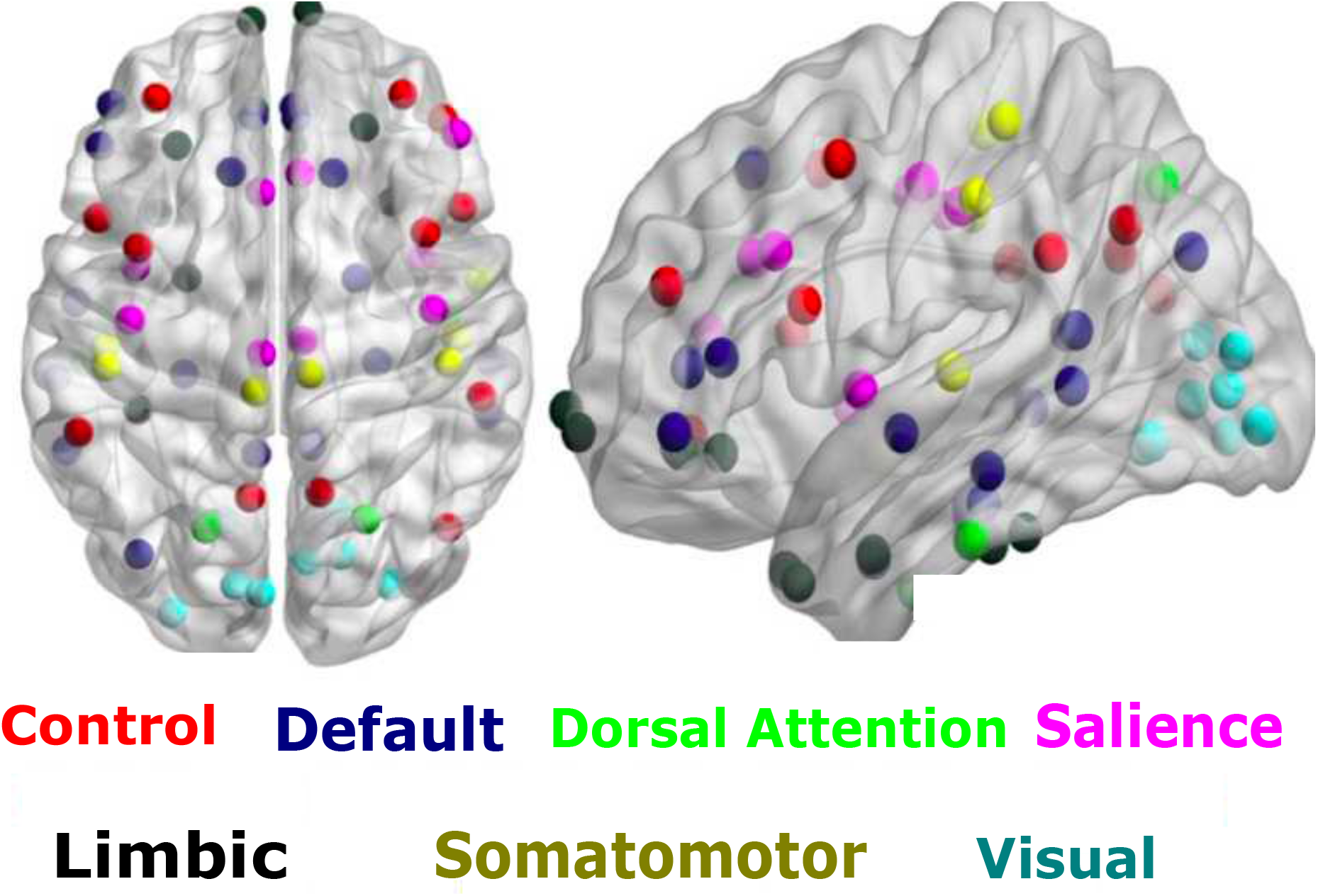
ROIs selected based on seven networks resting state parcellation proposed by Yeo et al. (2011) (cite). The ROIs (circles) related 3 prominent brain networks are overlaid on the spatial distribution maps derived from group ICA of multiple resting state networks of interest that is, the default mode network ROIs (DMN), the salience network ROIs (SN), and the Control network or Central Executive Network (CEN)

**Figure 3.**
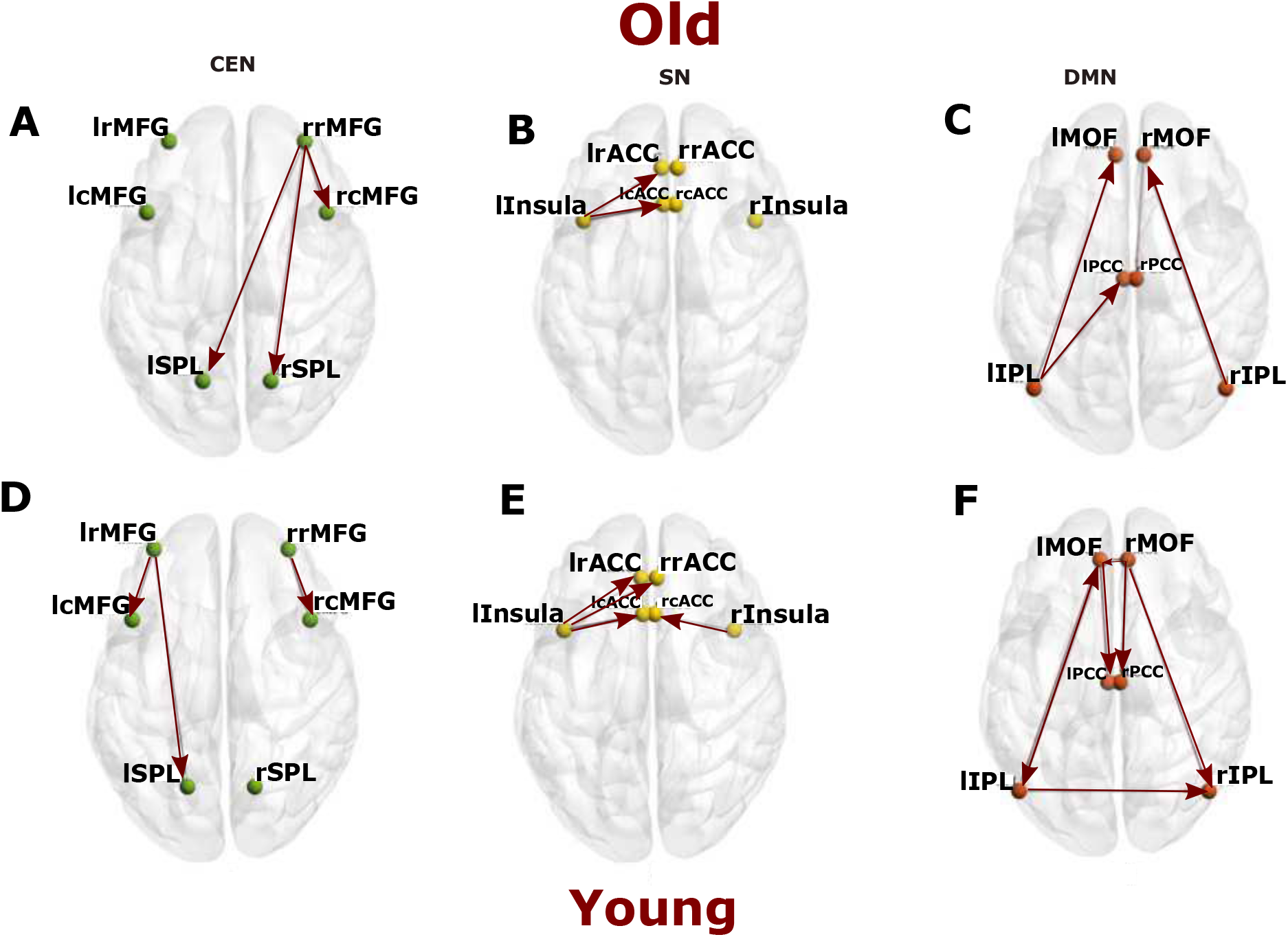
Directed Connectivity between the nodes of three resting state networks. Effective connectivity between **A.** six key nodes of CEN (green), **B.** six key nodes of SN (yellow), **C.** six key nodes of DMN (red) for elderly group. **D.** six key nodes of CEN (green), **E.** six key nodes of SN (yellow), **F.** six key nodes of DMN (red) for young group.

To further investigate the network properties, we quantified weighted net causal out-flow for the six nodes within CEN. The lRMFG and rRMFG (bilateral rostral areas in the Frontal Gyrus) acted as a causal outflow hub for young and elderly groups respectively. The majority of the identified causal outflows were significantly different for all the six nodes (p<0.01). With the exception of lRMFG and rRMFG, all nodes had negative causal outflow (causal inflow) for young individuals (**figure 4A**). In contrast, in the elderly group, other than rRMFG, only lCMFG had small positive outflow. All the remaining nodes of the CEN had causal inflow.

**Figure 4.**
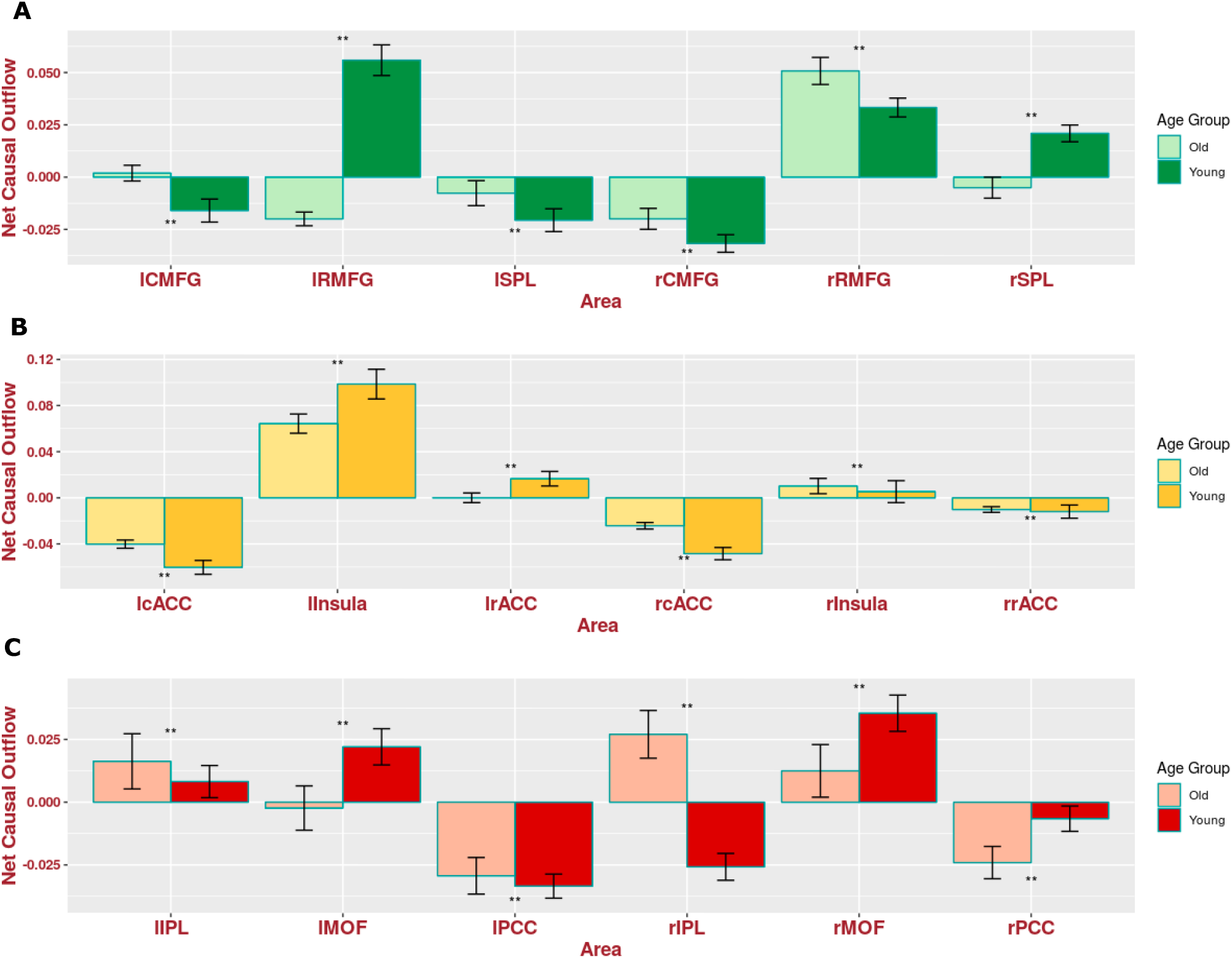
Weighted net causal flow for within network effective connectivity analysis. A. Weighted net causal outflow in nodes of the central executive network **B.** Weighted Net causal outflow in nodes of the salience network **C.** Weighted net causal outflow in nodes of default mode network. Weighted net causal outflows were significantly different in few nodes in each of the three RSN young and elderly group (p < 0.05 is indicated by ‘*’, p<0.01 is indicated by ‘**’, No significant difference is indicated by ‘NS’)

Next we proceed with analysis of the SN which comprise of six nodes (see **figure 2**), namely lInsula, rInsula, lcACC, rcACC, lrACC, rrACC. For within network analysis, quantitative comparison of MVGC analysis at the group level revealed a smaller number of significant directed causal influences compared with DMN and CEN. There was significant directed influence from lInsula to the lCACC, and the lRACC in young and elderly groups (**figure 3B, 3E**). Additionally, in the young group, significant directed influences were found from the rInsula to the rCACC and from the lRACC to the lInsula. One interesting find is that rInsula drives the rCACC in the young group but not in elderly group (**figure 3E**).

Weighted net causal outflow based on (out-In degree) analysis revealed lInsula as a major causal outlow hub in the SN for both age groups, but between group differences were also significant. In the elderly group, the causal outflow significantly reduced compared to young (Old < Young, p<0.01, FDR corrected). Furthermore, rInsula had a small causal outflow in both groups, and the group differences were statistically significant (p<0.01) and opposite of what is observed for the lInsula. The causal outflow from rInsula is significantly increased for the elderly group. lRACC had positive outflow for the young group and was significantly different for the elderly group (**figure 4B**). All the remaining nodes of ACC had negative outflow (causal inflow) for both groups. Causal outflows/inflows were significantly different (p<0.01) for all the nodes.

We next applied MVGC on the extracted time series for each of the six DMN nodes (see **figure 2**) for both young and elderly group to quantify the age effects in the dominant direction of influence (*p* < 0.05, false discovery rate (FDR) corrected). While in the younger group, GCA revealed significant directed causal connectivity from the lMOF (a driver as well as driven node) to the lIPL (bidirectional connectivity), lPCC (unidirectional) (follower nodes) and rMOF (a driver node) to the rIPL, rPCC, lMOF (interhemispheric directed connectivity), and lIPL to rIPL (interhemispheric connectivity) as shown in **figure 3F**, in elderly individuals, such bidirectional and interhemispheric causal connectivity was not present. Moreover, we see a reversal in the directed causal connectivity lIPL driving both lMOF and lPCC, and interhemispheric causal connectivity between lIPL to rIPL were completely absent (**figure 3C**) suggesting an age associated decrease in causal drive within DMN nodes.

Next, we estimated weighted net causal outflow or weighted (Out-In) degree in both young and elderly group. Based on the 100 bootstrap samples the distribution of weighted net causal outflow was calculated for each of the six nodes in DMN. For the young cohort, rMOF acted as a causal outflow hub among the nodes in DMN and was significantly different between the two groups ((Mann-Whitney U test p value < 0.001).), whereas rIPL acted as causal outflow hubhub for elderly group but acted as causal inflow hub for younger group (**figure 4C**). rPCC and lPCC both acted as causal inflow hubs receiving more drive from other nodes in the DMN for both young and elderly and were statistically significantly different in young group compared to old (Mann-Whitney test p value < 0.001). Interestingly, lPCC showed a reduction in causal inflow from young to old (Young > Old) while opposite was observed for the rPCC (Old > Young). Taken together, these results based on within network weighted causal flows (Out-In degree) suggest an age associated decrease in causal drive, and suggests decline in within network DMN functional connectivity.

### 3.2 Comparison of within network directed causal connectivity in young versus elderly using multivariate GCA in CEN, SN DMN with thalamo-cortical interactions

In elderly group, all the connections except rRMFG-rlSPL remained significant (*p* < 0.05, FDR corrected in the analysis including the thalamus. The connections between rRMFG-lSPL was mediated by the lThal. Some significant unidirectional causal connectivity was emerging from both the thalami, between lthal-lRMFG and rthal-lRMFG. In the CEN, we found frontal cortex was specifically driven by both left and the right thalamus in the left hemisphere. For both young and elderly groups, right and left thalamus was driving bilateral RMFG in the respective opposite hemispheres. Bilateral SPL was driven by rostral and caudal MFG in the elderly. Moreover, there were significant rostro-caudal interactions in the young as well as elderly group, but the direction of influence was reversed (**figure 5A and 5D**). For the young group, all the connections without thalamus were also found significant after inclusion of the thalamus (**figure 5**), which demonstrates that the overall patterns did not change by incorporating thalamo-cortial interactions. Rather, the evidence suggests that thalamic influence is absolutely critical to understand the variance of the prediction-errors for the estimation of directed connectivity in the triple resting networks. Additionally, the left thalamus exhibited directed connectivity among the following nodes of CEN: the lRMFG, the lCMFG, and the rRMFG (**figure 5D**). Right thalamus was the driver node and lRMFG was the follower node in the younger group.

**Figure 5.**
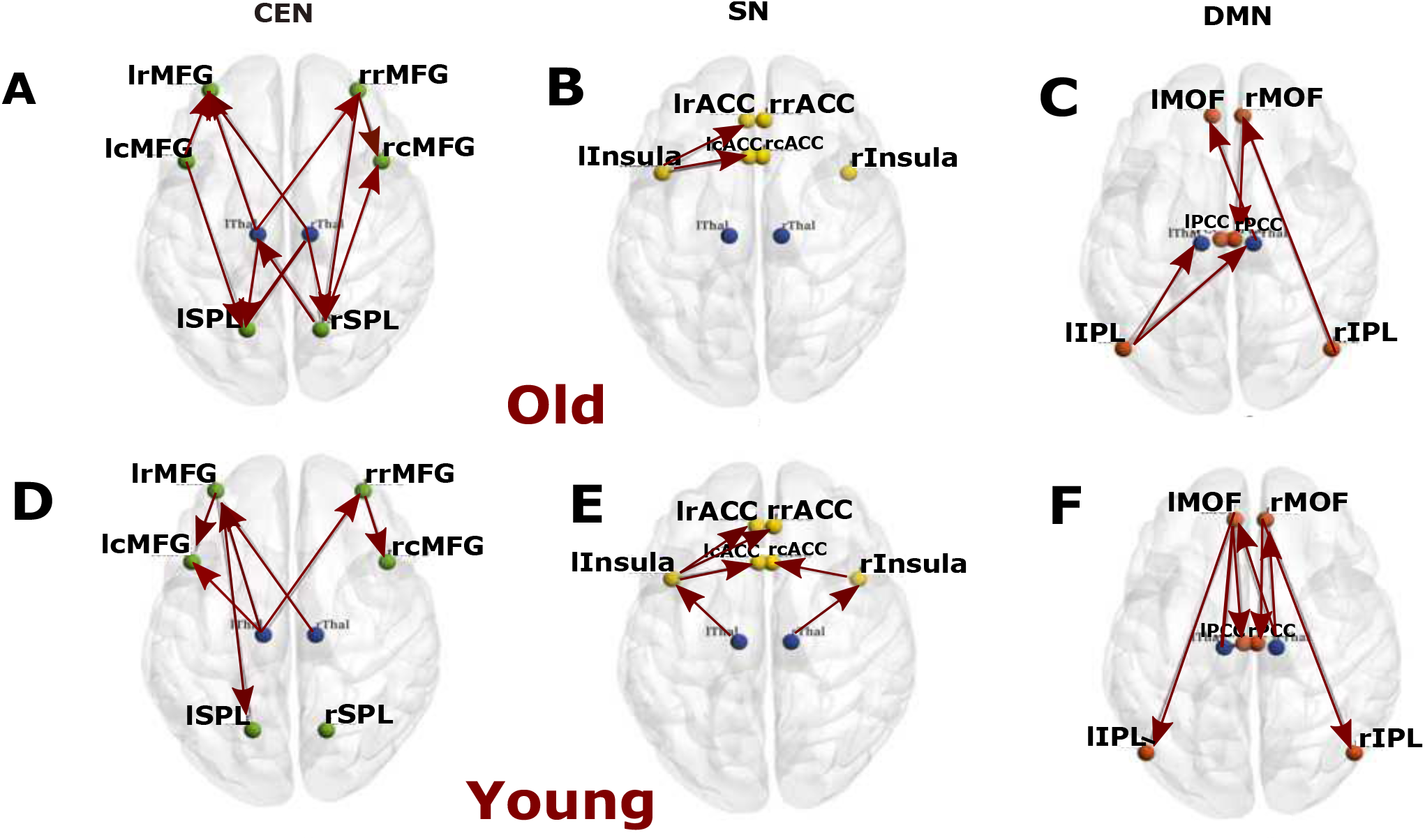
Effective Connectivity between the nodes of three resting state networks and the thalamus. **A.** Effective connectivity between six key nodes of CEN (green), **B.** six key nodes of SN (yellow), **C.** six key nodes of DMN (red) for elderly group in presence of thalamic nodes (blue). **D.** Effective connectivity between six key nodes of CEN (green), **E.** six key nodes of SN (yellow), **F.** six key nodes of DMN (red) for elderly group in presence of thalamic nodes (blue) for young group in presence of thalamic nodes (blue).

Net granger causal outflows were significantly changed after inclusion of thalamus for both the young and elderly groups. The effects of the thalamus were greater in the younger individual’s weighted (Out-In degree) net causal values (**figure 6A**). The left thalamus acted as a causal outflow hub for both the groups, as expected from the above results. However, the weighted causal outflow in the left thalamus was higher in the young group compared with the elderly group. We found significant group differences in weighted causal flow analysis for all the eight nodes (p<0.01).

**Figure 6.**
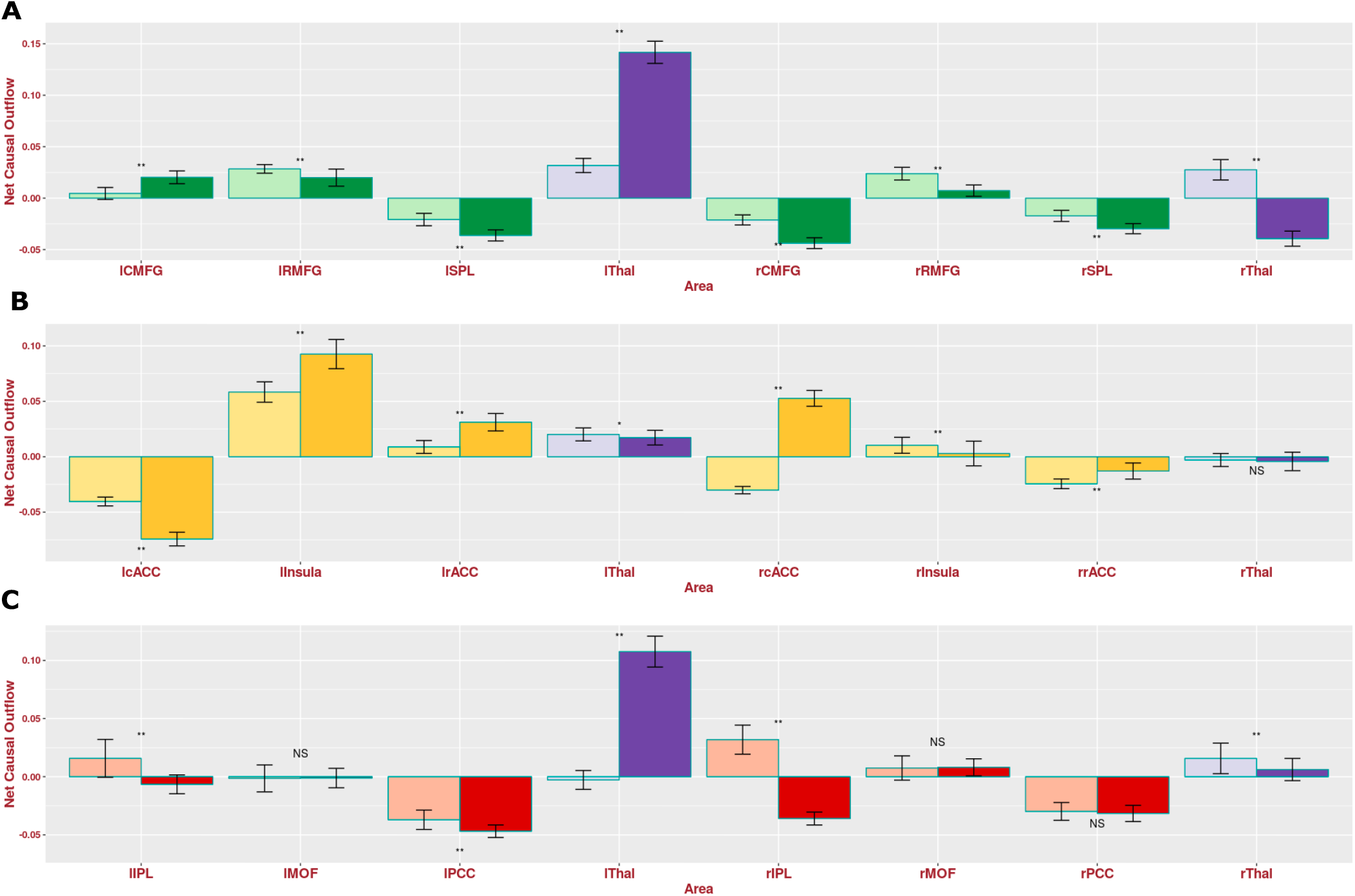
Weighted net causal outflow for within network effective connectivity analysis **A.** Weighted net causal outflow in nodes of the central executive network and two thalamic regions (left, right). **B.** Weighted net causal outflow in nodes of the salience network and two thalamic regions (left, right). **C.** Weighted Net causal outflow for default mode network and two thalamic regions (left, right). Weighted net causal outflows were significantly different in nodes in each of the three RSNs for young and elderly group (p < 0.05 is indicated by ‘*’, p<0.01 is indicated by ‘**’, No significant difference is indicated by ‘NS’).

Among the three resting state networks, the SN was least affected after inclusion of thalamus. No changes were found in causal structure after inclusion of thalamus in elderly individuals (**figure 5B**). In young individuals, significant causal connections were found from the lThal-lInsula connections (p < 0.01, FDR corrected) and from the rThal-rInsula connections (p < 0.01, FDR corrected) (**figure 5E**). No significant changes were found in the net causal outflow pattern in both the groups. Left thalamus exhibited positive outflow, higher in the case of older individuals (p<0.05). Right thalamus received marginally small negative outflow (inflow) for both groups (**figure 6B**).

Next, we studied causal interactions between thalamus and the DMN. We observed significant reconfiguration in the directed functional connectivity pattern found in the elderly individuals. On performing multivariate granger causal analysis on DMN after addition of thalami, some of the earlier significant causal connections disappeared while some thalamo-cortical causal connections emerged as important connections for both the age groups (**figure 5C, 5F**). The reversal of unidirectional connectivity between rIPL-rMOF follows a posterior-anterior gradient and anterior-posterior gradient, as seen previously in the connections rMOF-rIPL. Posterior-anterior directional connectivity continued to be the strongest in elderly group without taking into account thalamic interactions (**figure 5C**). Other than that, GCA revealed significant causal connections from the rMOF to the rPCC, from the rThal to the lIPl, from the lIPL to the rThal, from the rThal to the lMOF for elderly people (**figure 5C**). In the presence of thalamo-cortical causal interaction, directed functional connectivity between the left hemispheric nodes were largely absent in elderly. For the young group, the effect of the thalamus was more pronounced compared with the elderly group (Young > Old) (**figure 5F**). Instead of the connection from the lMOF to the lPCC, the connection from the lThal to the lPCC emerged as the strongest connection after the inclusion of the thalami (though the lMFCOF to the lPCC connection also remained significant). In addition, other significant connections were from the lMOF to the lIPL, from the lThal to the lMOF, from the lThal to rThal, from the lThal to the rPCC, from the rThal to the lMFC, from the rThal to the rMFC, from the rMFC to the rPCC, and from the rMFC to the rIPC (**figure 5F**), suggesting substantial effects of thalamus in reorganizing within network causal drive this network, and also revealing the effect is the strongest in the younger group compared with the elderly.

Net granger causal outflows were significantly changed after accounting for cortico-thalamic causal interactions, in particular, for the young group. Among the two thalami, the left thalamus emerged as a causal outflow hub for the young group, exhibiting substantial drive to the cortical nodes in the DMN Patterns in the weighted net causal outflows remained unchanged in elderly group (**figure 6C**). Right hemispheric IPL continued to be causal outflow hub for elderly group were significantly different (p< 0.01, FDR corrected) from the young group, even after accounting for thalamo-cortical interactions. Causal outflows were significantly different in both groups (p<0.01). Overall, we found stronger weighted causal outflow and increase causal drive from the thalamus among key nodes (left and right IPL, left and right PCC) of the DMN network associated with aging.

### 3.3 Reconfiguration of between network directed functional connectivity associated with age and cortico-thalamic interactions in triple networks

Next we asked in what specific way the between network interactions in the three neurocognitive networks are reconfigured by age and cortico-thalamic interactions. To address this systematically, we employed principal component analysis, (PCA) to combine the time series from each of the resting state networks (CEN, SN and DMN). We took the first principal component of all nodal time series of a network. Subsequently, we performed MVGCA among three nodes derived from the first principal component of each RSN. In between network analysis, SN exhibited stronger causal influence on both DMN and CEN (SN > DMN and SN > CEN) in both young and old groups (**figure 7A and 7B**). Directed functional connectivity and causal strengths were significantly stronger in old groups compared with young (*p* < 0.01, FDR corrected). CEN exhibited strong directed connectivity with DMN and hence, DMN was causally strongly driven by both SN and CEN (*p* < 0.01, FDR corrected). In general, we discovered the presence of stronger directed functional connectivity strength in the old group compared with the young group, suggesting an increase in between network causality with aging.

**Figure 7.**
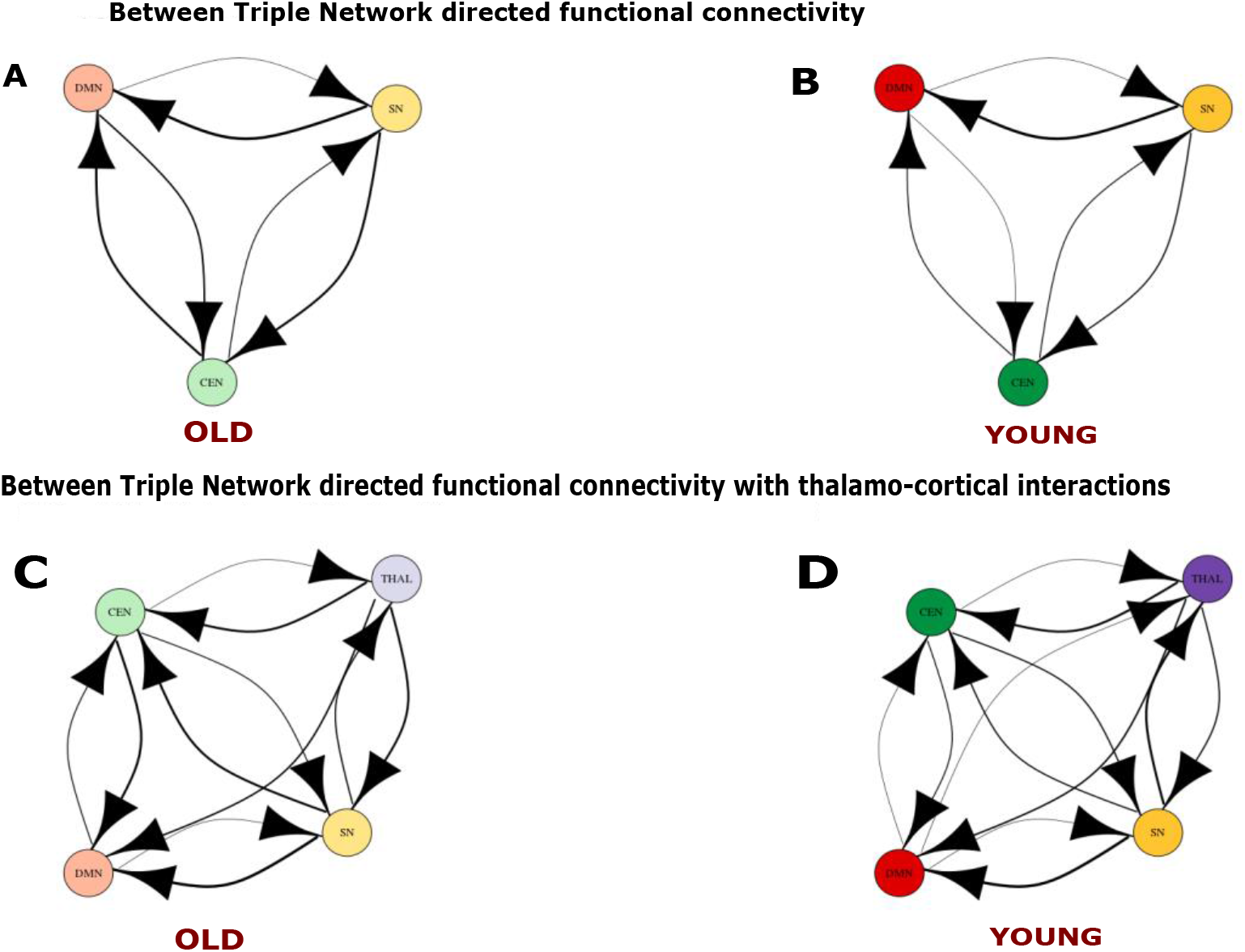
Effective connectivity between three resting state networks in absence and presence of the thalamus. **A.** Effective connectivity between three nodes representing three RSN, SN (yellow), DMN (red), CEN (green) for old population. **B.** Results for Effective connectivity between three nodes representing three RSN for young population **C.** Effective connectivity between three nodes representing three RSN and fourth node representing the thalamus (blue) for old population **D.** Effective connectivity between three nodes representing three RSN and fourth node representing the thalamus for young population

We then estimated the weighted net granger causal outflow in the triple networks Among the three RSNs, the SN was causal outflow hub. Network Causal outflows (In-out degree) were statistically significantly different between young and elderly groups (p< 0.01, FDR corrected) (**figure 8A**). Further, we repeated between network analysis, with the second principal component of all nodal time series of a network. No significant directed functional connectivity was found, confirming the fact that the first principal component sufficiently explained all the variabilities present in the time series of the three networks.

**Figure 8.**
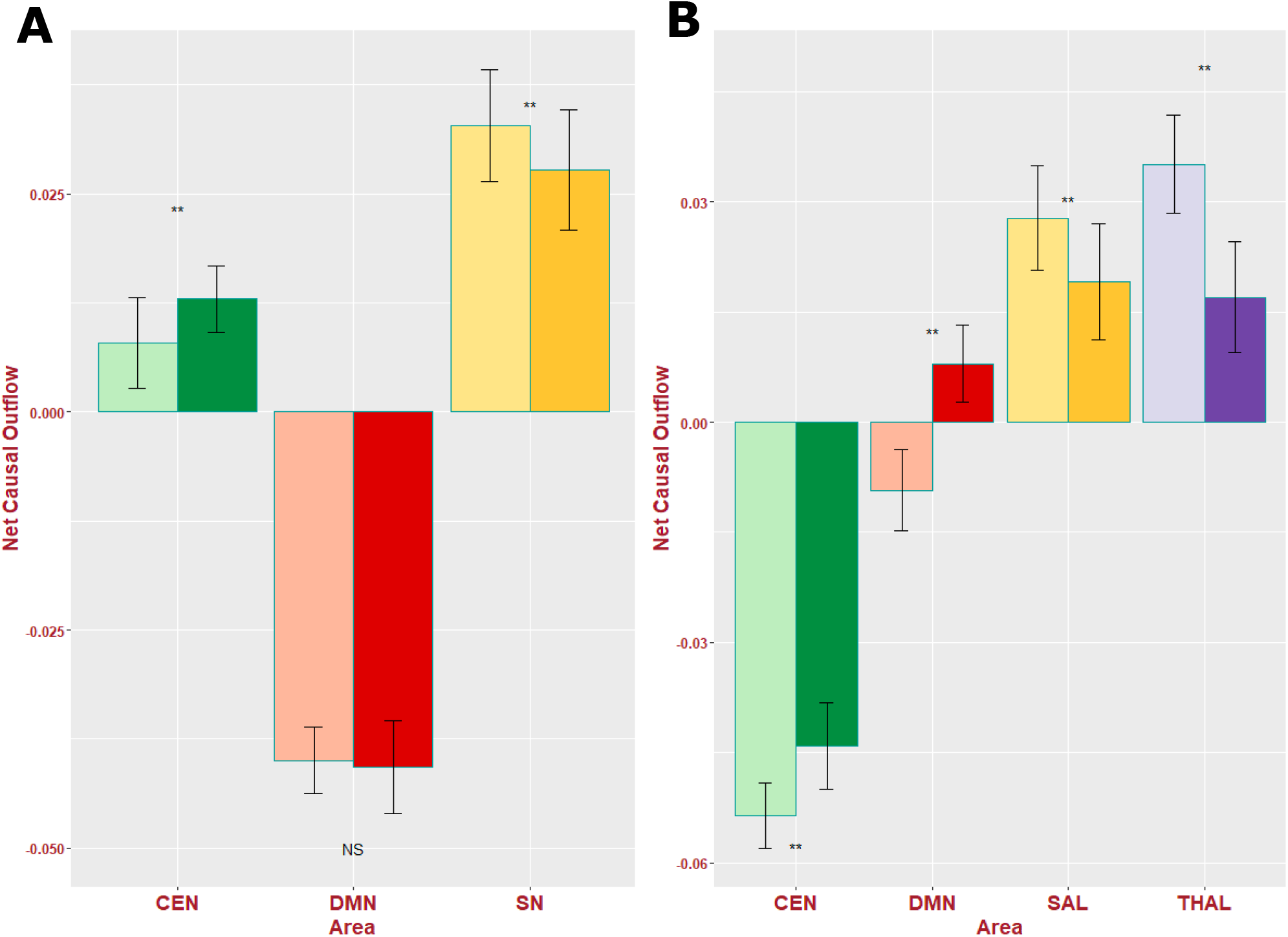
Weighted net causal outflow for between network effective connectivity analysis A. Weighted net causal outflow in three RSN **B.** Weighted net causal outflow in three RSN and the thalamus. Weighted net causal outflows were significantly different in few nodes in each of the three RSN young and elderly group (p<0.01 is indicated by ‘**’, No significant difference is indicated by ‘NS’).

Next we investigated between network causality in the presence of the thalamo-cortical interactions for both the groups. Directed functional connections from the thalamus to all three network nodes, namely the DMN, the SN, and the CEN in both age groups were found to be significant (*p* < 0.05, FDR corrected). The thalamus was also causally driven by CEN and SN for both elderly and young groups (**figure 7C and 7D**).

Weighted net causal outflows were significantly affected after taking into account thalamo-cortical interactions. We find evidence for the thalamus acting as a causal outflow hub (Old > Young) for the elderly group (**figure 8A**). Interestingly, SN and CEN dynamics were completely antagonistic with each other based on weighted causal outflow in the presence of cortico-thalamic interactions. DMN also exhibited greater causal inflow with respect to age. While SN received highest weighted causal outflow for young group (Young > Old, p<0.01, FDR corrected) CEN received highest causal inflow for elderly group (Old > Young) (**figure 8B**). Unlike the within network results, in between network analysis causal outflows were greater in the elderly group with or without accounting for thalamo-cortical causal interactions.

Overall, thalamo-cortical interactions did not necessarily alter the directed functional connectivity patterns and causality between the triple networks. After accounting for thalamo-cortical interactions, the causality dynamics between three resting state networks remained largely unaltered. However, the thalamus also received feedback causal influences and efferent drive from both the SN and CEN, and in turn the thalamus influenced resting state networks. Hence, there was strong evidence for bidirectional connectivity. Hence, the weighted net causal outflows for the elderly group were affected accounting for substantial thalamo-cortical interactions. However, this is hardly surprising given that left thalamus in particualr emerged as an important causal hub node in mediation of crucial thalamo-cortical interactions with respect to age. Taken together, within network thalamic drive progressively weakens with age, and stronger directed functional connectivity was found in the young group between thalamo-cortical connections. On the contrary, in the elderly group, between network directed functional connectivity was far less dissimilar accounting for thalamo-cortical interactions.

### 3.4 Replication Analysis

We identified a group of 24 young and 24 elderly participants from the publicly available Cambridge Aging Neuroscience dataset (https://camcan-archive.mrc-cbu.cam.ac.uk//dataaccess/) in the age range of 18–80 years who did not differ in mean age, gender distribution from Berlin dataset (see methods). Using this new dataset for independent validation, we conducted identical directed connectivity and weighted causal outflow analyses for each of the three-core neurocognitive resting state networks of interest.

In the replication analysis using the CAMCAN dataset, the six nodes for CEN included bilateral caudal middle frontal gyrus (rCMFG, lCMFG), rostral middle frontal gyrus (rRMFG, lRMFG) and superior parietal lobule (rSPL, lSPL). Multivariate Granger Causality was performed from the extracted time series to evaluate reconfiguration of within triple network directed functional connectivity in young and old age group.

We found several significant overlaps in the within network causality results between the connections between rRMFG-lSPL and lRMFG-rSPL and these connections were mediated by both the lThal and rThal respectively. Some significant unidirectional causal connectivity was emerging from both the thalami, between lthal-lRMFG and rthal-lRMFG ((p<0.05, FDR corrected). In the CEN, we found frontal cortex was specifically driven by both left and the right thalamus in the left hemisphere as was observed in the original data. For both young and elderly groups, right and left thalamus was driving bilateral RMFG in the respective opposite hemispheres. Bilateral SPL was driven by rostral and caudal MFG in the elderly. There were some differences as well. While there were significant rostro-caudal interactions as in the original data, but this time it was only present for the elderly and not for the young. There was interhemispheric directed connectivity between bilateral parietal lobule which was absent from the Berlin data. In replication analysis, we discovered in the elderly group rSPL (a driver node) was having significant directed functional connectivity (p<0.05, FDR corrected) with lSPL (a follower node). Significant connections (*p* < 0.05, FDR corrected) emerged from left and right thalami both in the young and old groups. However, the number of causal connections between thalamus and CEN nodes were greater in the younger cohort (**figure 9A, 9D**).

**Figure 9.**
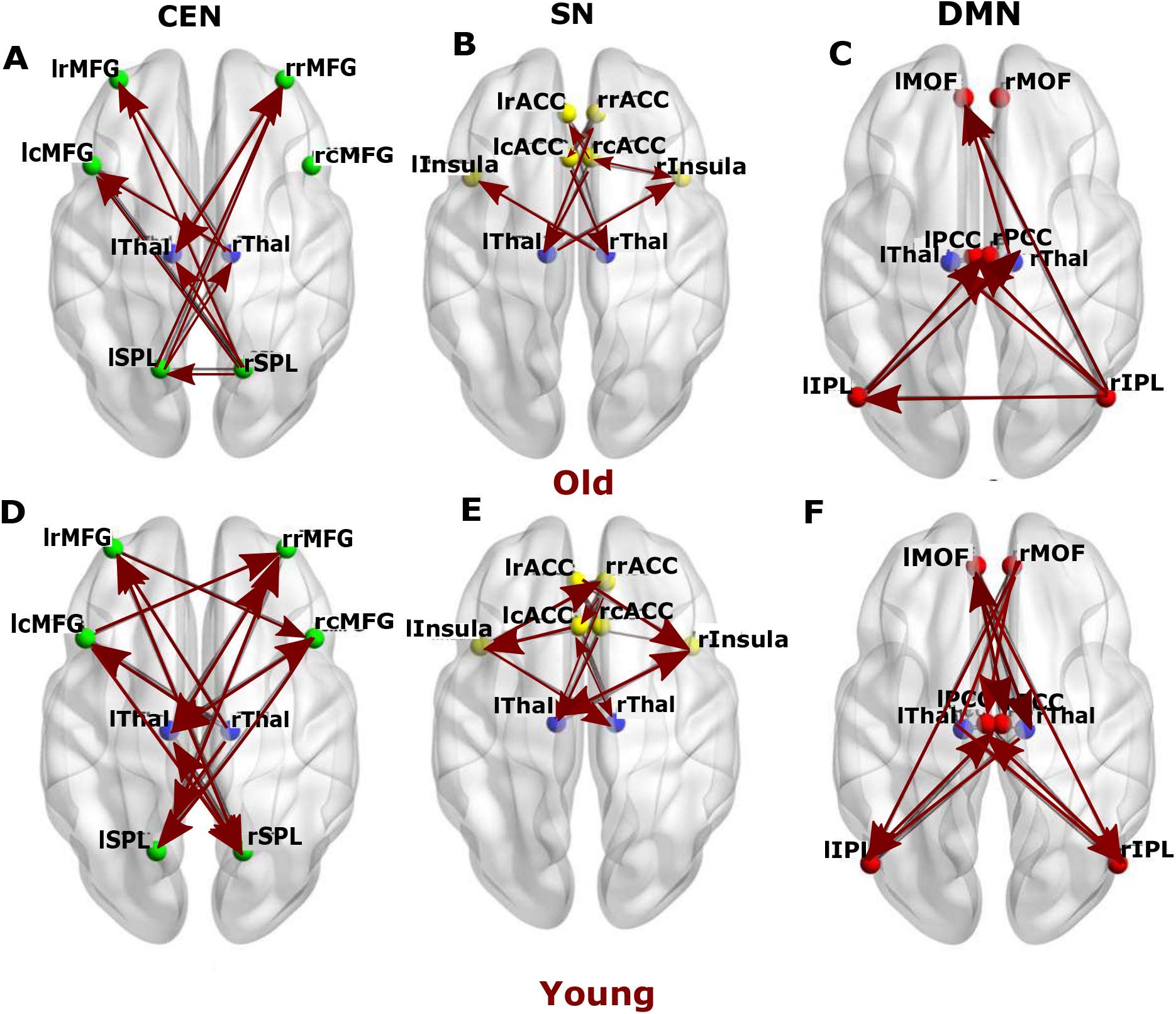
Directed functional Connectivity between the nodes of three resting state networks andthe thalamus in the replication analysis. Directed functional connectivity between **A**. key nodes of CEN (green), **B**. key nodes of SN (yellow), **C.** key nodes of DMN (red) for elderly group in presence of thalamic nodes (blue). **D.** Effective connectivity between key nodes of CEN (green), **E.** key nodes of SN (yellow), **f.** key nodes of DMN (red) for young group in presence of thalamic nodes (blue).

Bilateral rostral anterior cingulate (rrACC, lrACC), caudal anterior cingulate (rcACC, lcACC) and insula (rIns, lIns) were defined as the nodes of the SN. We observed directed functional connectivity between bilateral insula (driver) and caudal ACC (driven by Insula). We also found a difference while bilateral insula was driven by thalamus in the young group in the original data this connection was absent for the old group. In the replication analysis, this connection did not disappear in the old group (**figure 9B, 9E**). In both datasets, the number of connections decreases significantly in the old group compared to young.

In the DMN, the nodes selected were bilateral medial orbitofrontal (lMOF, rMOF), inferior parietal lobule (lIPL, rIPL) and Posterior Cingulate Cortex (lPCC, rPCC). In the younger group, GCA revealed extensive causal connections between all the DMN nodes (**figure 9F**) which was missing in the older group. In both data we found there is significant overlap between anterior-posterior interactions and the connections also exhibited hemispheric asymmetry in the older group.

The younger cohort exhibited a greater number of causal interactions than the older group (**figure 9C, 9F**), suggesting an age-related decrease in within DMN causal drive. Significant causal connections (*p* < 0.05, FDR corrected) emerged from left and right thalami both in the young and old groups. However, the number of causal connections between thalamus and DMN nodes was greater in the younger cohort (**Figure 9F**). We found also similarity in directed functional connectivity between bilateral IPL and bilateral MOF and missing lIPL-lMOF connections in original and replication analysis. In the replication data, we also discovered reversal of anterior-posterior directed connections between MOF and IPL associated with age as was found in the original data. There was a difference, rIPL was driven by rMOF in the original data and this directional functional connectivity reversed in older group. In the replication data, rIPL was driven by lMOF and not by rMOF and the directionality was similarly reversed in the older group as was found in the original data.

To represent the information of all the nodes in a particular network for a between network analysis, the time series of all the nodes in each of the resting state network were combined by employing a PCA (see methods). All causal connections were found significant. Between network directed functional connectivity showed significant overlap between SN, CEN, DMN connections in the young and old group in both data. In the young group, we discovered that SN (a driver network) exhibits strongest directional functional connectivity with both DMN (a follower network) and CEN (a follower network). Also, CEN drives DMN network causally and exhibits directed functional connectivity. In both replication data as wll as original data SN emerged as the key driver network exhibiting stronger directional functional connectivity with DMN and CEN associated with age. The number of directed connectivity as well strength increases with age which is an overlapping finding for both original and replication data (stronger between network causal connectivity in triple network interactions with age) (**Figure 10A, 10B**). Finally, thalamo-cortical interactions did not necessarily alter any of the directed functional connectivity patterns between the triple networks; however, displayed only directed connections only between CEN-Thalamus in the older cohort (**Figure 10C**). This was the main difference with original analysis where we discovered other connections such as Thalamus-SN, DMN-Thalamus and others. On the other hand, for both original and replication data in the young group we found presence of overwhelmingly large number of bidirectional functional connectivity of triple resting state networks with thalamus compared to elderly (**Figure 10D**).

**Figure 10.**
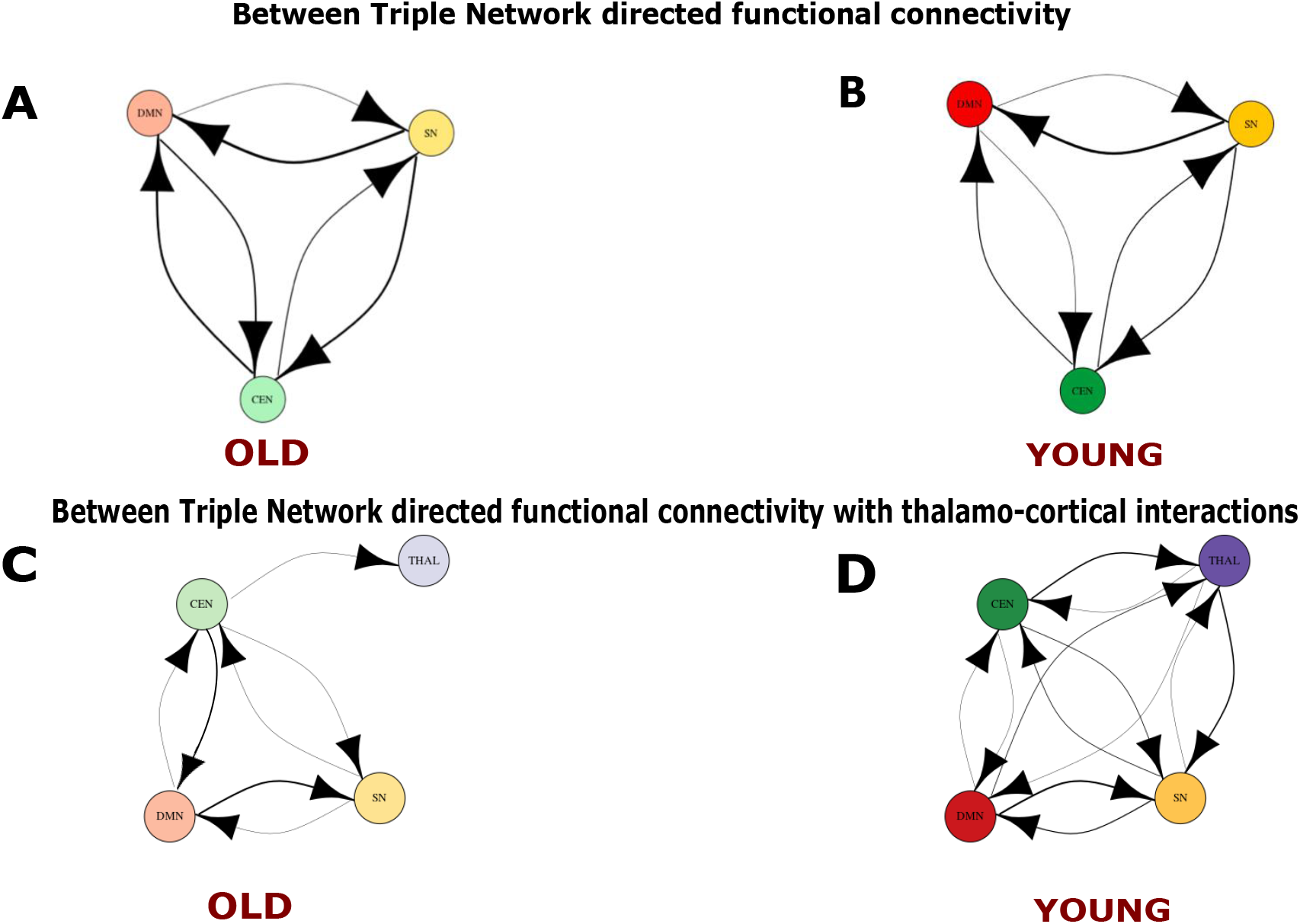
Directed functional connectivity between three resting state networks in absence and presence of the thalamus for the replication analysis. A. Directed connectivity between three nodes representing three RSN, SN (yellow), DMN (red), CEN (green) for old population. B. Results for Effective connectivity between three nodes representing three RSN for young population C. Directed functional connectivity between three nodes representing three RSN and fourth node representing the thalamus (blue) for old population D. Directed functional connectivity between three nodes representing three RSN and fourth node representing the thalamus for young population.

## 4. Discussion

Several recent studies have endeavored to unravel changes in the brain’s structural and functional connectivity with aging and the cognitive implications of these changes. The vast majority of these studies have used spatial and temporal correlation across different brain regions as measure of functional connectivity to characterize age-related changes in brain’s large-scale functional connectivity patterns (Vij et al., 2018). In the present study, we employ multi variate granger causality analysis to probe within- and between-network causal relationships among three key intrinsic resting state brain networks with the hope of facilitating more biologically meaningful interpretations of brain signatures of healthy aging.

Every cortical region receives feedforward projections from the thalamus and in turn sends outputs to one or multiple thalamic nuclei (Obeso et al.,2008); (McFarland et al.,2002). Thalamo-cortical projections relay nearly all incoming information to the cortex as well as mediate corticocortical communication (Sherman and Gui). Thus, deeper insight into brain functional characterization requires knowledge of the organization and properties of thalamo-cortical interactions. A recent study by ((Hwang, Bertolero, Liu, & D’Esposito, 2017)) showed the thalamus as an integrative hub for functional networks. A handful of studies also observed disrupted thalamic resting state functional networks in brain injury and schizophrenia ((Tang et al., 2011); (Wang, Rau, Li, Chen, & Yu, 2015a)). (Goldstone, Mayhew, Hale, Wilson, & Bagshaw, 2018) investigated the association of thalamic functional connectivity with behavioral performance in an elderly population. However, there is a knowledge gap in terms of how thalamo-cortical interactions sculpt causal drive and how they reconfigure directed connectivity in major neurocognitive networks with aging. This knowledge-gap is surprising given that lifespan-associated behavioral performance and flexibility in principle is governed by large scale functional brain network re-organization and whole brain dynamics at fast and slow time scales (Naik et al.; 2017, King et al.; 2018, Sahoo et al.; 2020).

Previous evidence suggests that there exist substantial subcortical-cortical causal interactions that are crucial for reconfiguration of brain dynamics in the triple networks with aging. Hence, focusing primarily on cortical nodes in these analyses paints an incomplete understanding. Here we investigated changes in causal dynamics among the triple networks accounting for thalamo-cortical interactions in our analysis for the first time. Overall, we found significant dynamic reconfiguration of between- and within-network directed functional connectivity and weighted causality change with aging. Our study also reveals the salience network’s role as a mediator of switching between DMN and CEN and establishes greater between network directional connectivity in older compared with younger adults. This finding is in line with the extant literature ((Menon & Uddin, 2010; Uddin, 2011); (Bonnelle et al., 2012)) and also provides confirmatory evidence of the pivotal role played by the SN for flexible switching as causal outflow hub. Finally, we also demonstrate how thalamo-cortical interactions play a crucial role in mediating within network interactions among the triple networks in the young group and how the causal drive gets diminished with age, suggesting age associated thalamo-cortical decline.

The question of segregated and integrated brain dynamics is fundamental and pertinent with regards to alteration and reconfiguration of brain network dynamics with aging. Hence, we focused on the decreased causal segregation of brain networks (i.e., increased internetwork directed connectivity), two features which can be considered a hallmark of the aging process. According to the dedifferentiation hypothesis of cognitive aging, age-related impairments in cognitive function arise from reduced distinctiveness of neural representations ((Li, Lindenberger, & Sikström, 2001)). Historically, the concept of dedifferentiation was introduced by Baltes and colleagues ((Baltes & et, 1980)) to account for age-related increases in the correlation between levels of performance on different cognitive tasks. At the neural level, numerous brain-imaging studies have shown that the aging brain adapts by exhibiting more global activation compared with younger individuals while performing a cognitive/motor task ((Cabeza, 2002); (Reuter-Lorenz, 2002);(Serrien, Ivry, & Swinnen, 2007); (Seidler et al., 2010)). In line with these finding, at the network level, several studies have found a decrease in within-network functional connectivity and an increase in between-network functional connectivity in RSNs with aging ((Andrews-Hanna et al., 2007);(Betzel et al., 2014);(Ferreira & Busatto, 2013);(Ferreira et al., 2016);(Geerligs et al., 2015);(Ng, Lo, Lim, Chee, & Zhou, 2016)). Overall, our study reproduces these results with directed functional connectivity analysis and further finds that the reorganization in causal connectivity patterns with age primarily reflects functional connectivity patterns. More specifically, younger individuals show increase in both the number and the strength of causal connections within DMN, CEN, and SN. At the between-network level, we find that causal strengths are significantly higher in the older individuals compared with the young, thus further substantiating the dedifferentiation hypothesis. Moreover, the present study also uncovers several novel observations. We observed the reversal of direction of causal connections between rMOF and rIPL with aging (change in directionality along anterior-posterior gradient), age-associated changes in the causal outflow in key nodes of triple networks, and reconfiguration of weighted causal outflow hubs. For future studies, it would be very interesting to see, by employing a similar methodology on a larger sample size including various stages of the adult lifespan, whether a clear trend emerges in directed connectivity patterns.

The role of the salience network in mediating switching between DMN and CEN is well established ((Sridharan et al., 2008); (Menon & Uddin, 2010); (Goulden et al., 2014)). In agreement with these observations, we find the SN to exert strong causal influence on both DMN and CEN in both age groups. In between network analysis, the SN is found to act as a causal outflow hub among the three RSNs we investigated. Interestingly, the causal influence of SN on both DMN and CEN increases with aging. An increase in the between network causality in the older group emerges as a general trend in our analysis. This is a counterintuitive result as within network analysis actually revealed significant evidence for thalamic decline with age. However, it seems higher neurocognitive networks such as SN, CEN, DMN establish stronger between network directed connections with the thalamus in the old group.

The thalamus has extensive connections with the entire cerebral cortex. By performing graph-theoretic analyses on thalamocortical functional connectivity data, ((Hwang et al., 2017)) demonstrated that the thalamus integrates multimodal information across diverse cortical functional networks and acts as an integrative hub for functional brain networks. The association between thalamo-cortical functional connectivity abnormalities and cognitive deficits in clinical conditions like schizophrenia ((Wang, Rau, Li, Chen, & Yu, 2015b),) is already known. Also known is the fact that thalamic volume significantly decreases with aging ((Walhovd et al., 2005); (Cherubini, Péran, Caltagirone, Sabatini, & Spalletta, 2009); (Zheng et al., 2018)). However, only a handful of studies (Goldstone et al., 2018) have investigated the thalamic influence in reorganizing RSNs with aging, and none have employed the directional connectivity analysis on triple network interactions with subcortical structures. In our analysis with directed functional connectivity measures, the thalamus emerges as an important node to critically influence both within- and between-network connectivity patterns among RSNs. After inclusion of the thalamus in the between network analysis, we observed that the thalamus acts as a causal outflow hub, driving all three network nodes in both age groups. In contrast, in within network analysis, the influence of the thalamus in reorganizing within network causality is much more prominent in the younger age group compared with the older group. In the DMN, the left thalamus comes out as a causal outflow hub for the young group, while patterns in the net causal outflows were unchanged in the older group compared with what was observed in the absence of thalamo-cortical interactions. While considering thalamo-cortical interactions, within network analysis in the CEN revealed that the left thalamus acts as a causal outflow hub for both the groups, however, the causal outflow strength was significantly greater in the younger compared with the older group. This preferential influence of the thalamus on younger individuals is consistent with the thalamic decline with aging, as discussed above.

Among the three resting state networks, the SN remains least affected after inclusion of the thalamus, both in within- and between-network analysis. We did not find substantial reconfiguration in SN with thalamo-cortical interactions in elderly individuals. Also, the re-configurations of this network were minimal in the young group. The net causal outflow pattern also did not change after taking thalamo-cortical interactions into account for both the groups. This could be related to the underlying decline in cortico-thalamic connectivity with aging. These findings are in concurrence with the observations made by the previous studies ((Wang et al., 2015b);(Cao et al., 2014); (Sakaki et al., 2016); (Xiao et al., 2018)) that in contrast to the DMN and CEN, within network connectivity is preserved or increased in SN with aging. Our result suggests that this preservation of causal connectivity patterns within SN may be crucial for aging and could be driven by thalamus.

We performed a replication analysis with an independent dataset controlling for age, gender, and some of the major findings with the original dataset were largely replicated. In within network analysis, a greater number of causal connections with higher strengths in the younger group compared with older individuals are consistent with our findings with the Berlin dataset. The effect of thalamo-cortical interactions was also revealed by greater directed connectivity from the thalamus to cortical nodes in the younger group compared with older individuals. Notably, we also found some specific differences between original and replication analysis. One of the ways we think this difference may have been attributed is the site variability, sample size, individual variability in the samples and data acquisition parameters. For example, an obvious difference is the duration of resting state scan and acquisition. They are certainly not identical in the two datasets (22 minutes for the Berlin data and 8 mins 40 secs for the CAMCAN replication cohort). This may actually contribute in creating differences in the directed connectivity patterns. However, what was reassuring is in spite of such clear differences the within and between network results between two samples were largely overlapping.

Finally, the thalamus is a heterogenous structure composed of several nuclei; each of which sends distinct afferent inputs to cortical regions as well as being driven by cortical outputs. Thus, probing the influence of different nuclei of thalamus on reorganization of within- and between-network causality of different RSNs would help to better describe the complex neurophysiological processes taking place in the brain with aging. The analysis presented here could be extended to other subcortical regions to understand how cortical-subcortical connectivity impact the cognitive performance and flexibility across age.

We acknowledge several limitations of our study that should be addressed in future work. The results presented here should be replicated in a larger lifespan cohort comprised of middle young and middle elderly groups to gain additional insights and true estimates of lifespan trends in causal outflow among core RSNs. Future research should also investigate the sensitivity of the analysis to the choice of different brain parcellation schemes. Moreover, application of Granger causality in fMRI data is not without limitations. Two potential deterrents are the fact that there is regional variability of the hemodynamic response to underlying neuronal activity, and the low sampling rate of fMRI makes it difficult to resolve this issue. MVGC detects causal interactions between brain regions by assessing the predictability of signal changes in one brain region based on the time course of responses in another brain region (Barnett et al. 2014; Stramaglia, S., Cortes, J. M., & Marinazzo, D. (2014); Seth and Barnett et al. 2018). Although there are some concerns that systematic differences across brain regions in hemodynamic lag can potentially lead to spurious estimations of connectivity (Rangaprakash, D. et al.; 2018), recent analyses also suggest that, when applied at the group level, GCA has good control over spurious results as age does not have a significant effect on the time course of the hemodynamic impulse response function or on the slope of the BOLD versus stimulus duration relationship. These results suggest that in cognitive studies of healthy aging, group differences in BOLD activation are likely due to age-related changes in cognitive–neural interactions and information processing rather than to impairments in neurovascular coupling (Grinband et al. 2017). Since the findings in the extant literature related to neurovascular couplings are mixed, we have further tested this by retrieving a proxy of the HRF at rest for both young and old group employing a recent pipeline (https://github.com/compneuro-da/rsHRF) (results of HRF retrieval and frequency domain analysis are not shown here to keep the findings less convoluted). Using the above measure and deconvolution on the unfiltered data we found practically no group difference in the retrieved HRF over the frequency ranges 0.01 Hz < f < 0.08 Hz. In particular, one might suspect that interregional differences in time-to-peak of the hemodynamic response function (HRF) function may lead to incorrect inference about the directed connectivity patterns among brain regions. Seth et al., (2013) and Barnett et al., (2014) however, established that since the HRF is effectively a slow-moving average filter, G-causality is principally invariant to it. However, ultrafast sampling is difficult to routinely employ in fMRI scans, and severe down sampling in fMRI still poses a serious challenge to G-causality analysis. On the other hand, the main strength of G-causality measure is that it can be applied directly to the given time series to detect the causal structure in any neuronal system without *a priori* hypotheses about the connectivity among the regions. The present study is one of the first to explore the reconfiguration of directed functional connectivity and causal dynamics in the triple networks in the presence of subcortical structures like the thalamus. Thus, we cannot restrict ourselves with a particular hypothesis. Moreover, the prime focus of the present study is to identify the changes in G-causality among the RSNs with ageing and with the inclusion of subcortical structures, rather than to identify the ground truth G-causality patterns. The information obtained from the present study could be used to implement more hypothesis driven characterization of the networks in the future, such as that offered by Dynamic Causal Modelling (DCM) or its spectral variant (REF). It is also worth noting that several neuroimaging studies have exploited this exploratory nature of Granger causality to estimate directed functional connectivity for both electrophysiological and BOLD signals (REFs).

In conclusion, the results of the present study demonstrate that directed functional connectivity and weighted causal analysis can provide critical insights regarding within- and be-tween-network information flow across the lifespan over and above insights already provided by existing functional connectivity studies. This study also establishes the thalamus as a com-mon driver of organization and reconfiguration of triple network dynamics with aging, a conclusion that should encourage future research to explore the influences exerted by subcortical structures on cortical networks and their cognitive and clinical implications.

## Acknowledgements

This study was supported by NBRC Core funds, Ramalingaswami Fellowships (Department of Biotechnology, Government of India) to DR (BT/RLF/Re-entry/07/2014). AB also acknowledges the support of the Centre of Excellence in Epilepsy and MEG. DR was also supported by SR/CSRI/21/2016 extramural grant from the Department of Science and Technology (DST) Ministry of Science and Technology, Government of India. AB and DR are also supported by BT/MED-III/NBRC/Flagship/Program/2019. LU is supported by R01 MH107549 grant from National Institute of Mental Health (NIMH) USA. Replication Data used in this study was generously provided as a public resource and repository by the Cambridge Centre for Ageing and Neuroscience (CamCAN). CamCAN funding was provided by the UK Biotechnology and Biological Sciences Research Council (grant number BB/H008217/1), together with support from the UK Medical Research Council and University of Cambridge, UK.

